# Individual differences in visual search performance extend from artificial arrays to naturalistic environments

**DOI:** 10.1101/2021.10.15.464609

**Authors:** Thomas L. Botch, Brenda D. Garcia, Yeo Bi Choi, Caroline E. Robertson

## Abstract

Visual search is a universal human activity in naturalistic environments. Traditionally, visual search is investigated under tightly controlled conditions, where head-restricted participants locate a minimalistic target in a cluttered array presented on a computer screen. Do classic findings of visual search extend to naturalistic settings, where participants actively explore complex, real-world scenes? Here, we leverage advances in virtual reality (VR) technology to relate individual differences in classic visual search paradigms to naturalistic search behavior. In a naturalistic visual search task, participants looked for an object within their environment via a combination of head-turns and eye-movements using a head-mounted display. Then, in a classic visual search task, participants searched for a target within a simple array of colored letters using only eye-movements. We tested how set size, a property known to limit visual search within computer displays, predicts the efficiency of search behavior inside immersive, real-world scenes that vary in levels of visual clutter. We found that participants’ search performance was impacted by the level of visual clutter within real-world scenes. Critically, we also observed that individual differences in vi^1–3^sual search efficiency in classic search predicted efficiency in real-world search, but only when the comparison was limited to the forward-facing field of view for real-world search. These results demonstrate that set size is a reliable predictor of individual performance across computer-based and active, real-world visual search behavior.

## Introduction

Locating an object in a cluttered environment is a ubiquitous visual behavior. The mechanisms by humans accomplish visual search have been comprehensively studied in both artificial arrays (Treisman & Gelade, 1980) and complex scene images (Wolfe, Võ, et al., 2011). Yet, little is known about whether the principles of visual search revealed by these studies extend to naturalistic visual contexts, where real-world environments are actively explored from a first-person perspective.

One key factor known to limit search performance is set size: the number of items within a visual array. Increasing set size impairs search performance in both artificial arrays (Neider & Zelinsky, 2008; Palmer, 1994) and pictures of complex scenes (Castelhano & Henderson, 2007; Henderson et al., 2009). However, it remains unclear whether set size effects analogously limit behavioral performance during active exploration of real-world environments, where environmental structure and memory are available to aid attentional guidance (Bar 2004, Vo 2019). Further, to our knowledge, whether individual differences in search efficiency in artificial displays predict naturalistic search performance in real-world environments has never been explored. This knowledge gap has important implications for understanding clinical conditions such as autism (Plaisted et al., 1998) and informing real-world applications such as radiology (Wolfe, 2020), where computer-based visual search paradigms are often used to model real-world behavior.

Here, we leverage advances in virtual reality (VR) technology to relate individual differences in classic visual search paradigms to naturalistic search behavior. Participants completed two tasks: (1) a classic conjunctive search paradigm with arrays varying in set size and (2) a naturalistic search behavior inside of immersive, real-world environments varying in levels of visual clutter (Rosenholtz et al., 2007). Across the two tasks, we characterized the impact of set size on visual search performance. We also tested whether efficiency was related across artificial and naturalistic contexts.

## Methods

### Participants

25 adults participated in two experiments (N=16 females; mean age 22.96 +/-4.08 STD years). Participants were recruited based on (1) having normal or corrected-to-normal vision and no colorblindness, (2) having no neurological or psychiatric conditions, and (3) having no history of epilepsy. Written consent was obtained in accordance with the Declaration of Helsinki via a protocol approved by the Dartmouth College Committee for the Protection of Human Subjects (CPHS).

### Remote data collection

Participants received a standalone headmounted display (Oculus Quest 2, www.oculus.com, single fast-switch LCD, 1832×1920px per eye; ∼90° field of view; 72 Hz refresh rate) preconfigured with the ManageXR (www.managexr.com) device management software. Experiments were built in Unity version 2018.4.12f1 (www.unity.com) with custom scripts written in C#. Experimental data was collected through a custom pipeline written in C# and PHP. Specifically, a virtual private server (VPS) was created though Amazon Web Services Lightsail and configured with the Linux, Apache, MySQL, and PHP (LAMP) stack. An additional Dropbox SDK (www.github.com/kunalvarma05/dropbox-php-sdk) was used to connect lab Dropbox accounts to the VPS allowing for direct transmission of data from the HMD to Dropbox.

### Experiment 1 – Naturalistic Visual Search

#### Exp. 1 – Stimuli and set size manipulation

In the naturalistic search experiment, stimuli consisted of 360° “photospheres” of real-world scenes, sourced from an online photo sharing website (www.flickr.com). We curated 54 photospheres with four criteria to minimize the complications of defining set size in real scenes (Wolfe, Alvarez, et al., 2011). First, we selected photospheres of indoor scenes, as outdoor scenes contain few segmented regions which may not be representative of the true set size. Second, we ensured the photospheres did not contain humans to avoid the possibility that humans are a unique object category. Third, we confirmed that each photosphere contained a “singleton” target object: an object that appeared only once inside a given photosphere. Fourth, given the importance of distance to scene processing in early visual areas on the brain (Kravitz et al., 2011), we ensured that all photospheres had comparable depth. To this end, we estimated the depth of each photosphere using the big-to-small (BTS) algorithm (Lee et al., 2020).

We adopted the concept of visual clutter as a proxy for set size in real-world scenes (Neider & Zelinsky, 2008; Rosenholtz et al., 2007) and approximated the visual clutter of each photosphere using the proto-object segmentation algorithm (Yu et al., 2014). Subsequently, we divided the photospheres into three bins (18 photospheres each) based on the estimated clutter measurements (low, medium, and high clutter) and ensured that the average clutter of each bin significantly differed from the others (Figure 1A). The average depth of photospheres in each quantile did not significantly differ (Figure 1B).

**Figure 1.**
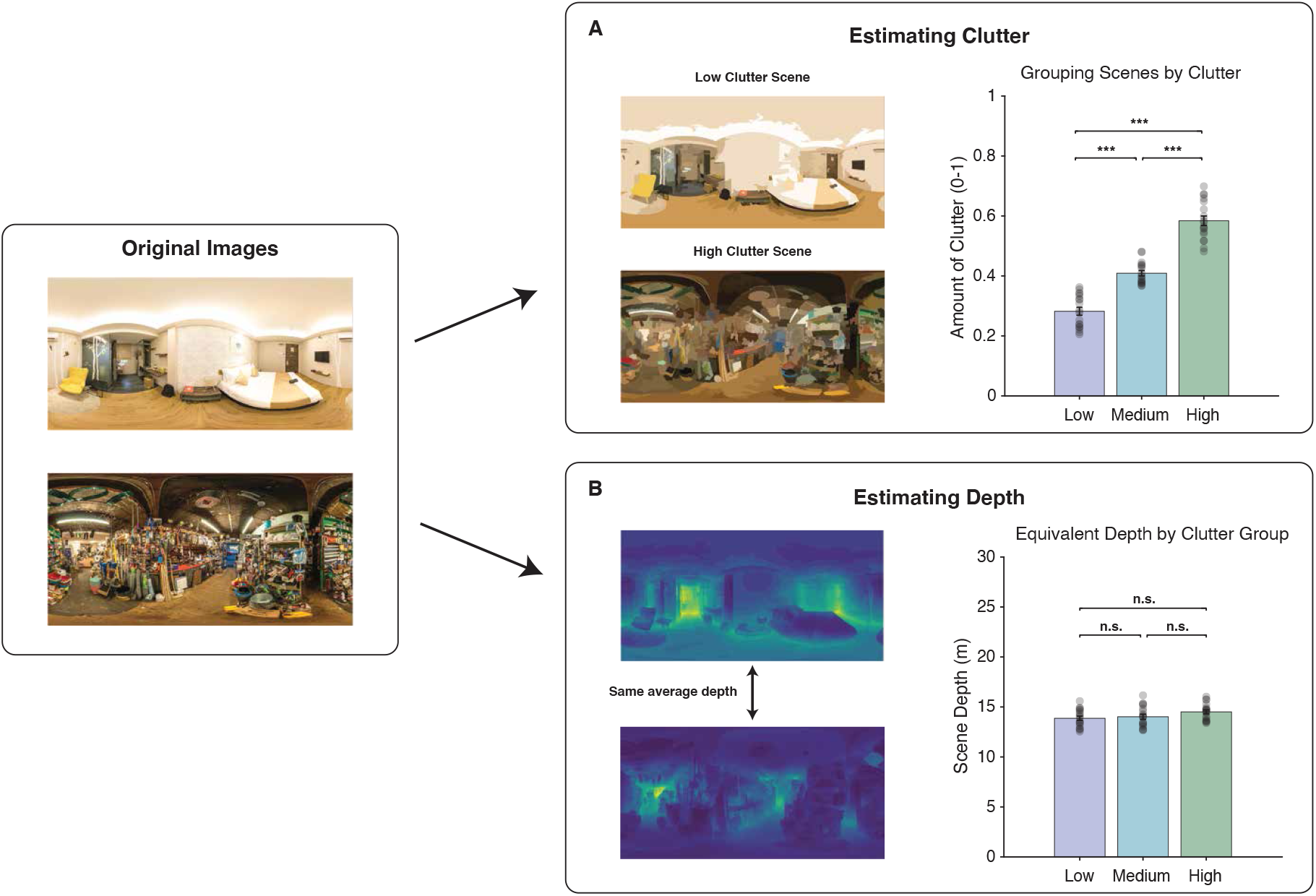
Schematic of visual clutter and depth estimation. **(A)** Example visualizations of visual clutter estimated by the proto-object segmentation algorithm. Photospheres were divided into three quantiles, and average clutter of each quantile significantly differed from the others (*F*_(2,51)_ = 144.7, *p* < 0.001). **(B)** Example visualizations of scene depth estimated by the big-to-small algorithm. The average depth of each clutter quantile did not significantly differ (*F*_(2,51)_ = 1.20, *p* = 0.31). In all plots, error bars represent 1 SEM. \**p* < 0.05, \*\**p* < 0.01, \*\*\**p* < 0.001, n.s. *p* > 0.05 difference between conditions.

Target object locations were balanced across photospheres within each clutter bin. For each scene, the yaw of each photosphere was randomly rotated such that the target object was located in one of three quadrants of the immersive environment relative to the participant’s initial facing direction: (1) to the left of the participant, (2) in front of the participant, or (3) to the right of the participant. This resulted in an equal distribution of target object locations relative to the participant across the three possible quadrants (6 photospheres per quadrants), and across the clutter bins (18 photosphere per quadrant).

#### Exp. 1 – Paradigm

On each trial of the naturalistic visual search experiment (54 trials), participants were presented with a photosphere via the headmounted display (HMD) for a maximum of 30 seconds, or until the controller trigger was pressed indicating detection of the target object (Figure 2A). In all scenes, an occluding wall obstructed the 90° immediately behind the participant such that the 270° in front of the participant was visible. Accordingly, participants were informed that the area behind them would not be visible and instructed to explore the forward, left, and right portions of the photosphere. To mitigate confusion during the real-world visual search task, we informed participants that the target object would always be present inside the virtual environment.

**Figure 2.**
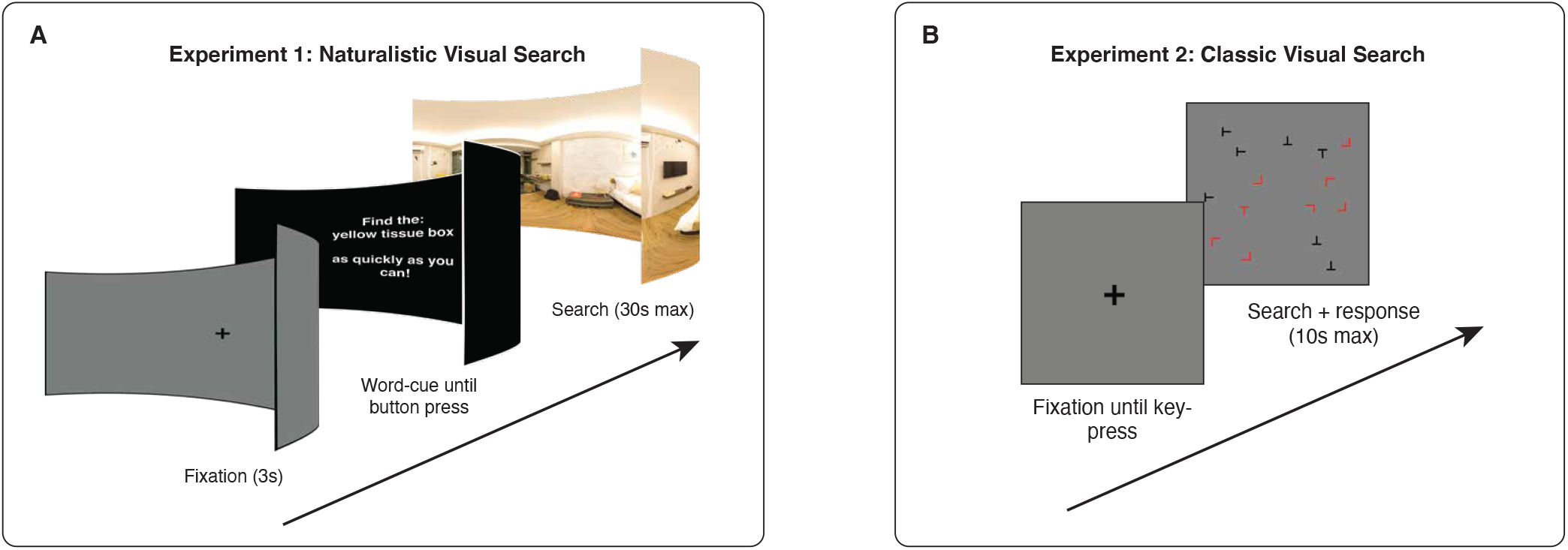
Experimental paradigms. **(A)** Naturalistic visual search paradigm. After a pre-trial fixation, participants were presented with a conjunctive word description of a target object. Participants subsequently searched for the described object inside a photosphere. **(B)** Classic visual search paradigm. After a pre-trial fixation, participants searched for a red T embedded within a cluttered array.

Before each trial, participants were presented with a pretrial fixation target at screen center to ensure participants entered each photosphere facing the same direction. Participants were required to align their head-center with the target for 3 seconds. Subsequently, participants were presented with a conjunctive word cue (e.g. green bottle) describing the target object in the following photosphere. Participants were instructed to “find the target as quickly as possible”. To report the target, participants centered their head on the target (specifically, they centered a light gray circle, which was locked to screen center, on the target) and pressed the controller trigger. A response was considered correct if the participant’s head coordinate was within a 7.5° visual angle radius from target center when the trigger was pressed, and reaction time was calculated as the time of the trigger press relative to trial start. After pressing the trigger, participants were given feedback on the accuracy of their response. The gray, head-locked circle would turn green if the participant selected the correct object and would turn red if the participant selected an incorrect object. After each trial, participants were returned to a virtual home environment where they were informed of their reaction time and instructed to take a break. A mandatory break occurred after each quarter of the experiment (14 trials) to allow participants to rest their eyes.

At the start of the study, participants were shown a set of instructions orienting them to the task. Following the instructions, participants completed two practice trials to ensure familiarity with the task. Participants were highly accurate during practice trials (mean accuracy: 92%), indicating comprehension of the task.

### Experiment 2 – Classic Visual Search

#### Exp 2 – Stimuli and set size manipulation

In the classic visual search experiment, stimuli consisted of letter arrays, which were presented on a grey background around a central fixation point (Figure 2B). The letters in the array had two feature dimensions: form (Ts and Ls) and color (red and black). Arrays spanned 25°x25° visual angle, and letters within the array were randomly distributed around a central fixation point and spaced from others by 2° visual angle. Displays had three potential set size conditions: 5, 15, or 25 letters.

#### Exp 2 – Paradigm

On each trial of the classic conjunctive search task (180 trials), participants were instructed to report the presence/absence of a target letter (a red T) using a keypad. Note, the target letter shared a feature dimension with each type of distractor (black Ts and red Ls). There were two trial types, target present or target absent, which each occurred 50% of the time. On trials without a conjunction target, an additional distractor was added at random.

Each trial lasted for a maximum of 10 seconds or until a key press. Before each trial, participants were shown a black fixation cross and required to press a button to start the trial. Participants were instructed to fixate on the cross until trial start, after which point they were free to move their eyes. Participants were instructed to “find the target as quickly as possible” and to “press 4 if the target is present or 6 if the target is absent”. Participant reaction time was calculated as the time of the button press relative to trial start. Following each trial, participants were given feedback on the accuracy of their response (a green check for correct responses and a red X for incorrect responses). A mandatory break occurred every 45 trials to allow participants to rest their eyes.

At the start of the study, participants were shown a set of instructions orienting them to the task. Following the instructions, participants completed a set of practice trials (12 trials) to ensure familiarity with the task. Participants were highly accurate during practice trials (mean accuracy: 100%), indicating comprehension of the task.

### Motor response control task

To acclimatize participants to selecting targets in VR and to establish a base-rate reaction time for each participant, a motor response control task was presented at the beginning of each session. On each trial (36 trials), participants were presented with a gray photosphere containing a red dot (target) within their front-facing field of view at a random location around a circle with a radius of either: 7.5°, 17.5°, or 27.5° visual angle from world center (12 trials each, balanced for visual quadrant) (Supplementary Figure 1A). Participants were instructed to move a gray circle, locked to the center of the HMD, to the red dot and press the controller trigger as quickly as possible. There was a non-significant difference in reaction time as stimuli moved further from world center (Supplementary Figure 1B; *F*_(2,72)_ = 1.94, *p* = 0.151).

## Results

To investigate if classic findings of visual search extend to naturalistic settings, we developed a novel paradigm in which participants locate objects inside 360° real-world scenes. In the naturalistic visual search task, we evaluated the impact of visual clutter, an estimate of set size for scenes, on visual search performance. In the classic visual search task, we estimated the impact of set size effects on individual performance using a minimalistic visual display of letters. Finally, we directly compared individual efficiency, the slope of the relationship between set size and reaction time, within each task to understand the relationship between computer-based measurements of visual search and naturalistic search behavior.

### Naturalistic Visual Search Performance

We first examined the relationship between visual clutter and search performance inside immersive, real-world scenes. Overall, we found that participants were faster and more accurate to locate the target in lower-as compared with higher-cluttered scenes. Combining data across our participants, we found a significant correlation between clutter-level and reaction times to correctly detect a target (Figure 3A; *r*_*s*_ = 0.654, *p <* 0.001). This correlation was significant in all three sections of the environment (right, left, and center of the participant) (left frame: *r*_*s*_ = 0.709, *p* < 0.001; front frame: *r*_*s*_ = 0.806, *p* < 0.001; right frame: *r*_*s*_ = 0.536, *p* = 0.024). A one-way ANOVA revealed a significant main effect of clutter on reaction times across scenes (Figure 3B; *F*_(2,51)_ = 12.74, p < 0.001). This significant effect of clutter is further demonstrated in the individual participant average reaction times for each clutter quantile (Figure 3C; *F*_(2,72)_ = 76.25, *p* < 0.001) along with individual participant false alarm rate (ANOVA: *F*_(2,72)_ = 11.09, *p* < 0.001). Overall, these results suggest that visual clutter modulates visual search performance inside real-world scenes.

**Figure 3.**
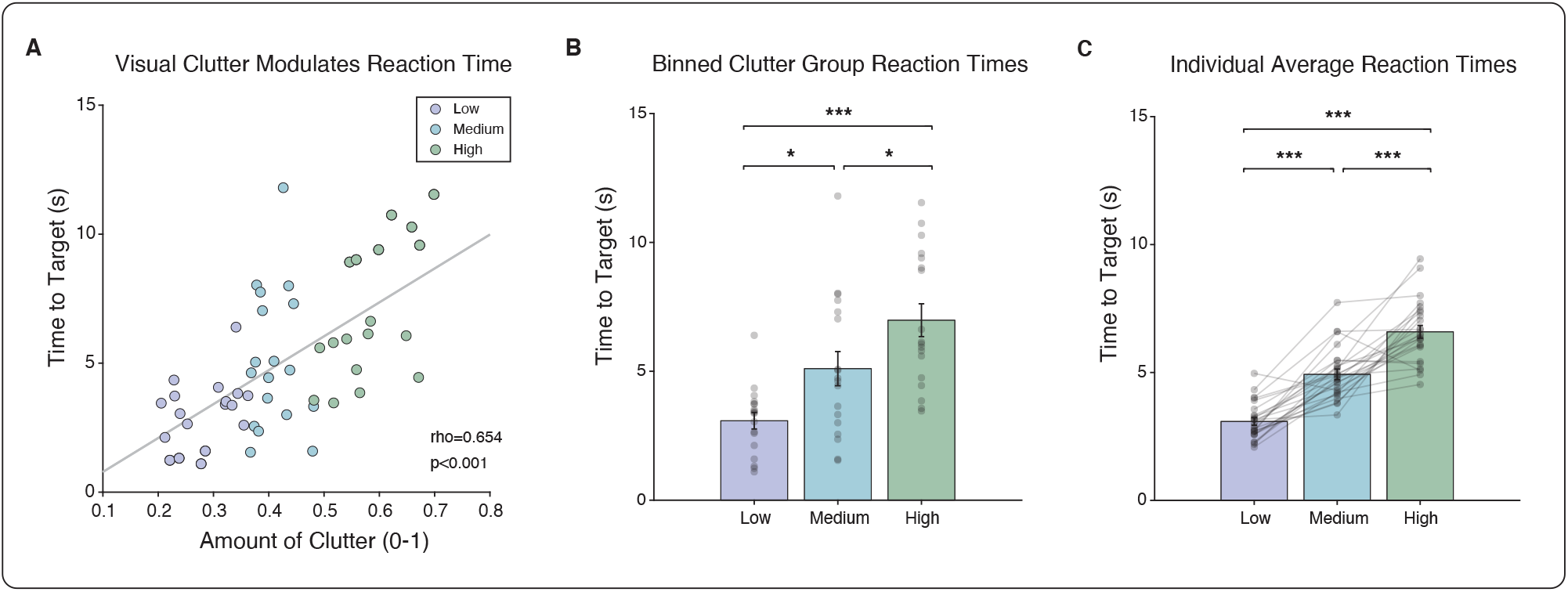
Experiment 1 - Naturalistic visual search results. **(A)** Amount of visual clutter correlates with reaction time in real-world environments (*r*_*s*_ = 0.654, *p* < 0.001). **(B)** Average reaction time across participants for each scene binned into clutter groups (*F*_(2,51)_ = 12.74, *p* < 0.001). **(C)** Individual participant average reaction times binned into clutter groups (*F*_(2,72)_ = 76.25, *p* < 0.001). In all plots, error bars represent 1 SEM. \**p* < 0.05, \*\**p* < 0.01, \*\*\**p* < 0.001, n.s. *p* > 0.05 difference between conditions.

### Classic Visual Search Performance

We next evaluated the relationship between set size and search performance in a classic visual search paradigm. Within each set size, we calculated the average reaction time of each participant separately for the target present and target absent trials. For target present trials, we performed a one-way ANOVA which revealed a significant main effect of set size (Figure 4A; *F*_(2,72)_ = 62.81, *p* < 0.001). Similarly, we found a main effect of set size for target absent trials (Figure 4B; *F*_(2,72)_ = 52.41, *p* < 0.001). Finally, we calculated the false alarm rate of each participant within each set size and performed a one-way ANOVA which showed no main effect of clutter (*F*_(2,72)_ = 1.77, *p* = 0.177). In sum, these results dovetail with previous findings of classic visual search paradigms demonstrating the impact of set size on visual search performance (Wolfe & Horowitz, 2017).

**Figure 4.**
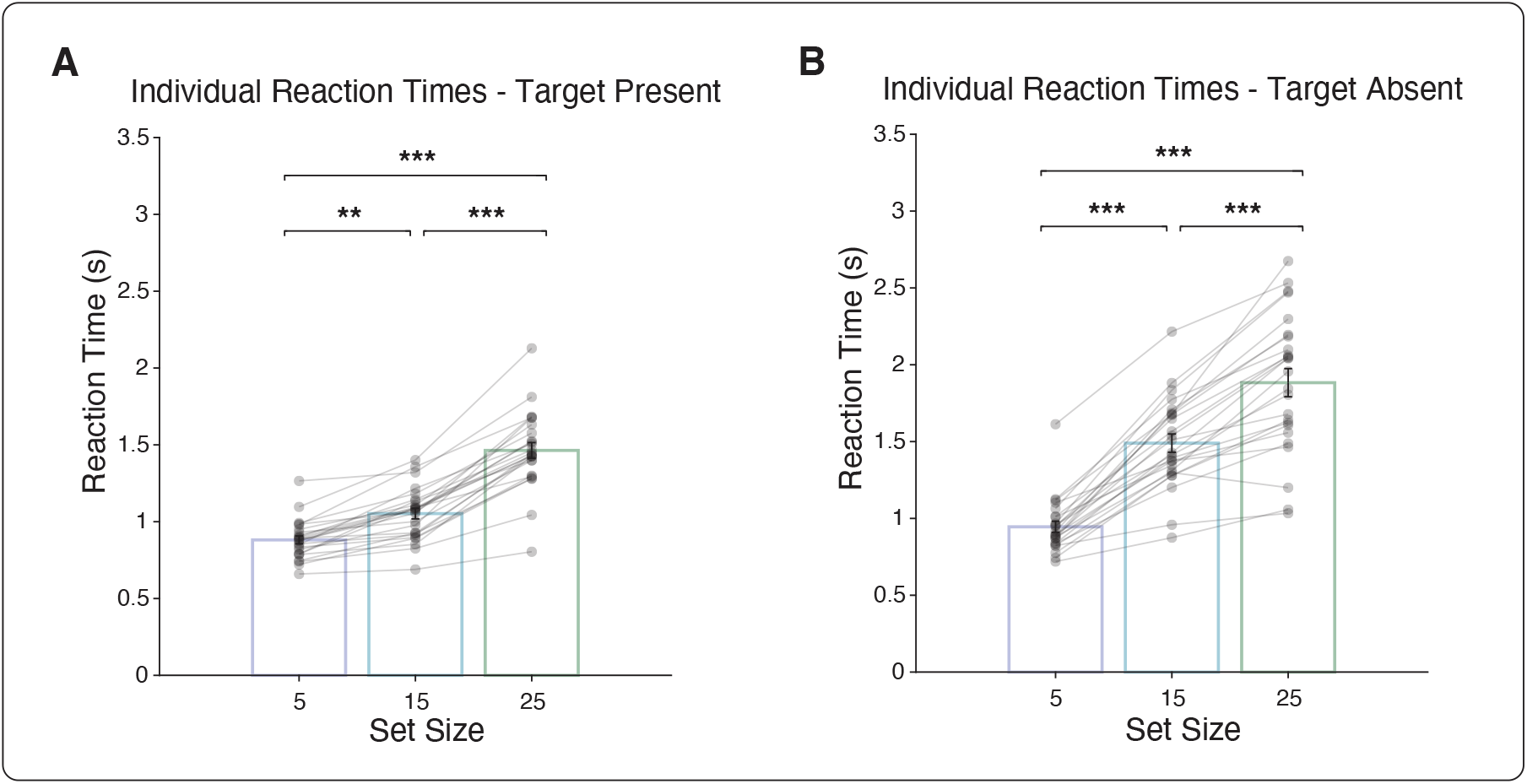
Experiment 2 -Classic visual search results. **(A)** Individual reaction times are modulated by set size on target present trials (*F*_(2,72)_ = 62.81, *p* < 0.001) and **(B)** target absent trials (*F*_(2,72)_ = 52.41, *p* < 0.001). In all plots, error bars represent 1 SEM. \**p* < 0.05, \*\**p* < 0.01, \*\*\**p* < 0.001, n.s. *p* > 0.05 difference between conditions.

### Reliability of Individual Reaction Times

Before examining the relationship between the two experimental paradigms, we established the reliability of individual reaction times for each task by iteratively calculating split-half reliability. Specifically, within each task for each participant, we divided reaction times within each set size in half and calculated the average Spearman’s correlation over 10,000 unique combinations. For the naturalistic visual search task, we conducted a one-sample t-test which showed that participant reaction times were significantly reliable (*t*(24) = 10.92, *p* < 0.001). Likewise, for the classic visual search task, we performed a one-sample t-test which revealed that participant reaction times were significantly reliable (*t*(24) = 8.17, *p* < 0.001).

### Relating Performance on Naturalistic and Classic Visual Search Tasks

Having established that each experimental paradigm exhibits within-individual reliability, we next investigated how individual differences in search performance on the two tasks related to one another. Within each task, we calculated the z-scored efficiency, the slope of a linear function mapped between set size and reaction time, of visual search for each participant. We first compared individual efficiency in the front quadrant of the naturalistic visual search task with each trial type of the classic visual search task. We found a significant relationship between search efficiency on the naturalistic search task and the target present trials of the classic visual search task (Figure 5A; *r*_*s*_ = 0.428, *p* = 0.034). The correlation between the naturalistic search and classic search tasks remained significant even when regressing out motor response times obtained from the motor response control task (*r*_*s*_ = 0.426, *p* = 0.038), suggesting that this finding was not mediated by individual differences in motor response times. We found a similar relationship of individual efficiency between the front frame of the naturalistic search task and the target absent trials of the classic visual search task (Figure 5B; *r*_*s*_ = 0.537, *p* = 0.006), which again remained significant even when regressing out motor response times obtained from the motor response control task (*r*_*s*_ = 0.537, *p* = 0.007). Together, these results suggest that efficiency on a classic visual search task, indexed by a set size manipulation, predicts efficiency in naturalistic visual search, indexed by a clutter manipulation in complex, visual scenes.

**Figure 5.**
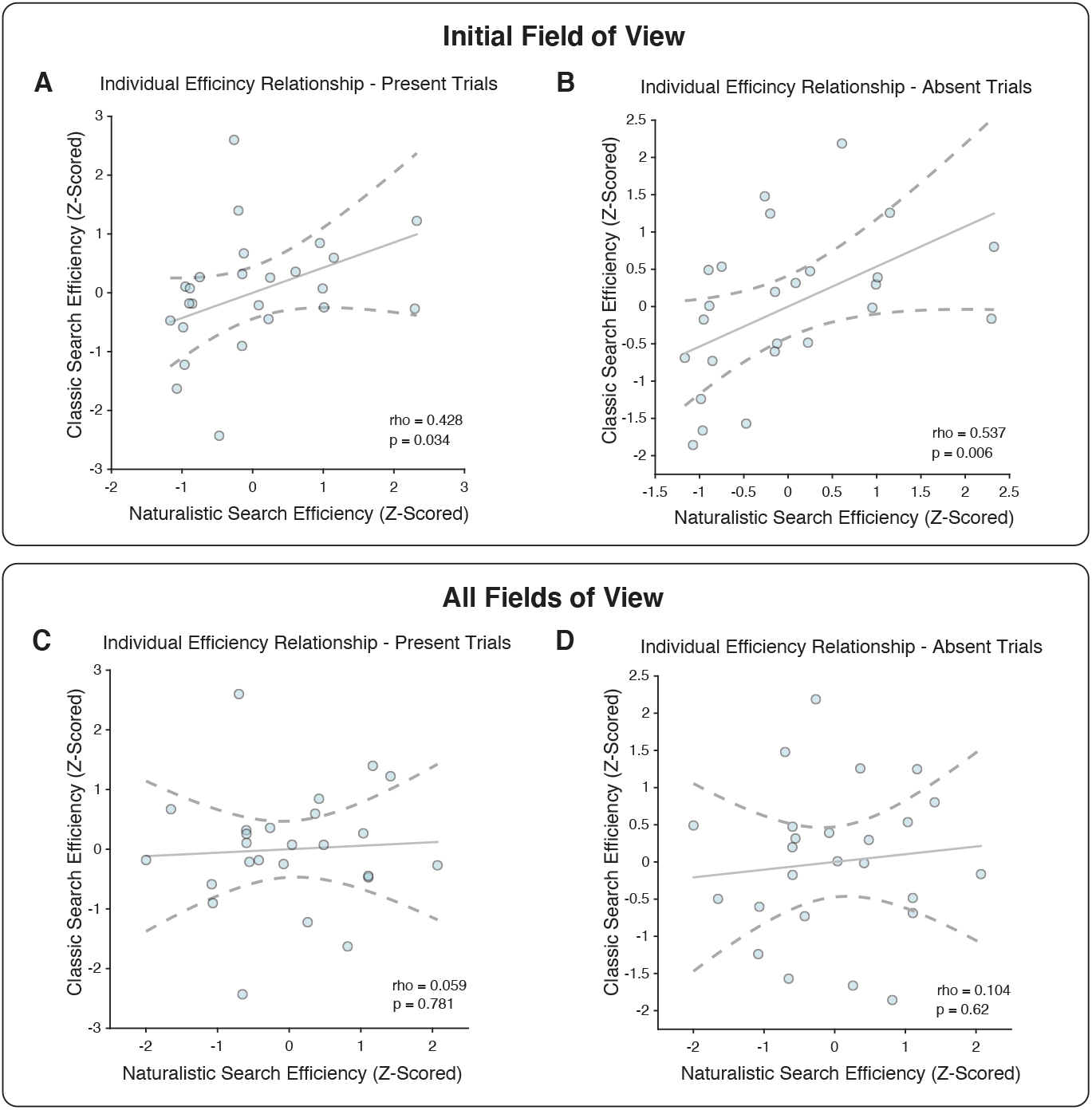
Relating individual performance across tasks. An individual’s efficiency on the classic visual search task predicts efficiency within the initial field of view of the naturalistic visual search task for both **(A)** target present (*r*_*s*_ = 0.428, *p* = 0.034) and **(B)** target absent trials (*r*_*s*_ = 0.537, *p* = 0.006). **(C**,**D)** However, when all fields of view are considered in the naturalistic visual search task, there is no longer a relationship between the two tasks. This occurred for both **(C)** target present (*r*_*s*_ = 0.059, *p* = 0.781) and **(D)** target absent trials (*r*_*s*_ = 0.104, *p* = 0.62).

Importantly, this relationship between naturalistic and classic search performance did not hold when left and right quadrants were included in the analysis (Figure 5C; *r*_*s*_ = 0.059, *p* = 0.781). Likewise, we found no relationship between individual efficiency in the naturalistic visual search task and the target absent trials of the classic visual search task (Figure 5D; *r*_*s*_ = 0.104, *p* = 0.62). Further, we found no relationship between the left and right frames of the naturalistic visual search task to the target present and absent trials of the classic visual search task (left-present: *r*_*s*_ = 0.022, *p* = 0.917; left-absent: *r*_*s*_ = -0.017, *p* = 0.937; right-present: *r*_*s*_ = -0.021, *p* = 0.922; right-absent: *r*_*s*_ = -0.039, *p* = 0.853). These results suggest that the increased motor and working memory demands associated with searching through an immersive environment to out of sight locations may disrupt the relationship observed between classic and naturalistic search.

## Discussion

We find that visual search in immersive, real-world environments bears remarkable similarities to classic search in two important senses. First, naturalistic and classic search respect a common principle of “set size efficiency”: just as classic search efficiency is limited by the “set size” of a visual display (i.e. the number of distractors in the display), so naturalistic search efficiency is limited by a real-world analogue of “set size”, visual clutter (i.e. the number of non-target objects in a real-world environment). Second, individual differences in classic and naturalistic search efficiency are related: individuals with steeper costs of set size in a classic search arrays were more severely hindered by visual clutter in real-world environments, although only when targets appeared in the initial field of view. Together, these findings suggest that classic search is an excellent model of the pervasive real-world activity of visual search and suggest that action contributes an additional layer of complexity to behavior.

Naturalistic paradigms present a valuable opportunity to validate and extend models of human behavior derived in laboratory settings to the conditions and demands of everyday life (Felsen & Dan, 2005; Leopold & Park, 2020). Within the vision sciences, advances in virtual reality technology present a promising avenue to investigate behavior within naturalistic contexts while simultaneously balancing experimental control (Doucet et al., 2016; Scarfe & Glennerster, 2015). This opportunity enables researchers to exact experimental rigor while presenting diverse real-world stimuli and compare models of visual processing within active visual settings (Haskins et al., 2020). Accordingly, an increasing number of studies employ virtual reality headsets to investigate visual function under naturalistic constraints (Beitner et al., 2021; Cohen et al., 2020; Li et al., 2018), providing essential connections between computer-based findings and naturalistic behavior. Yet, few studies have sought to relate models of visual performance on classic paradigms with behavior in real-world settings.

Most previous studies of naturalistic search have primarily used computer-based paradigms featuring photographs of real-world scenes. Such studies have revealed many attributes guiding attention in real-world scenes not present in classic search (Wolfe, Võ, et al., 2011). For example, recent studies of search using naturalistic photographs reveal the important interplay of memory (Võ & Wolfe, 2012), semantic structure (Vo & Henderson, 2009; Võ et al., 2019), and top-down attentional guidance (Castelhano & Heaven, 2010; Castelhano & Henderson, 2007) during the visual search of scenes. Based on these findings, it is not a given that basic principles of classic search, such as the relationship between set size and visual search efficiency, would extend to naturalistic settings where additional cues guide search behavior.

Recently, a few studies have investigated search performance in head-mounted displays using virtual environments. These studies have found that active exploration of and interaction with environments bolsters memory and aids visual search performance (Draschkow & Võ, 2017; Li et al., 2018). Additionally, a recent study found that engaging spatial priors through perceptual priming increases search efficiency (Beitner et al., 2021), further underscoring a pivotal role of memory in visual search. While these findings uncover important top-down properties guiding attention, the degree to which basic features limit visual search performance in real-world scenes is left unknown.

To our knowledge, no previous study has, to our knowledge investigated the common mechanisms underlying classic and active, naturalistic search. Our results therefore build on previous work in two important ways. First, we demonstrate that set size, a property that limits visual search, continues to limit performance during naturalistic search behavior in photospheres of real-world scenes. Second, we show that while an individual’s search efficiency is correlated between the two visual search tasks within the initial field of view, the incorporation of action alters the relationship of behaviors. All in all, our findings dovetail with previous studies in suggesting that naturalistic behavior engages additional mechanisms that guide attention during visual search.

Certainly, our experimental paradigm has shortcomings. First, in contrast to many studies of visual search in which eye-tracking measures are employed, we were only able to use a combination of head-tracking data and key-press reaction times. This method is undoubtably noisier than measuring eye-tracking reaction times in each task. However, given the close coupling of head and eye movements (Freedman, 2008) and the presence of set size effects within both paradigms, we do not believe a different measurement would drastically alter our results. Second, dissimilarities of the stimuli used in each paradigm, specifically the minimalistic nature of letter displays as opposed to real-world scenes, creates difficulties in drawing a direct comparison across the two tasks. That said, the correlation of individual efficiency between the two paradigms within the initial field of view makes it unlikely that stimulus content hindered the direct comparison of tasks. A continuum of stimulus naturalism exists, moving from well-controlled psychophysics displays into real-world settings, which may expose divergent guiding mechanisms of attention dependent on stimulus content. Thus, future studies may look to alter the stimulus along this naturalistic continuum; for example, performing searches for isolated objects on complex, real-world backgrounds.

In sum, we find that visual clutter limits visual search performance in immersive, real-world scenes. Individual efficiency relates across a naturalistic and classic visual search task when considering targets within participants’ immediate field of view, but this connection disappears when active exploration was required to find the target. Together, these findings suggest that active search alters the properties guiding attention during visual search and highlight the importance of relating computer-based and naturalistic tasks to better inform models of behavior.

## Supplemental material

**Supplementary Figure 1.**
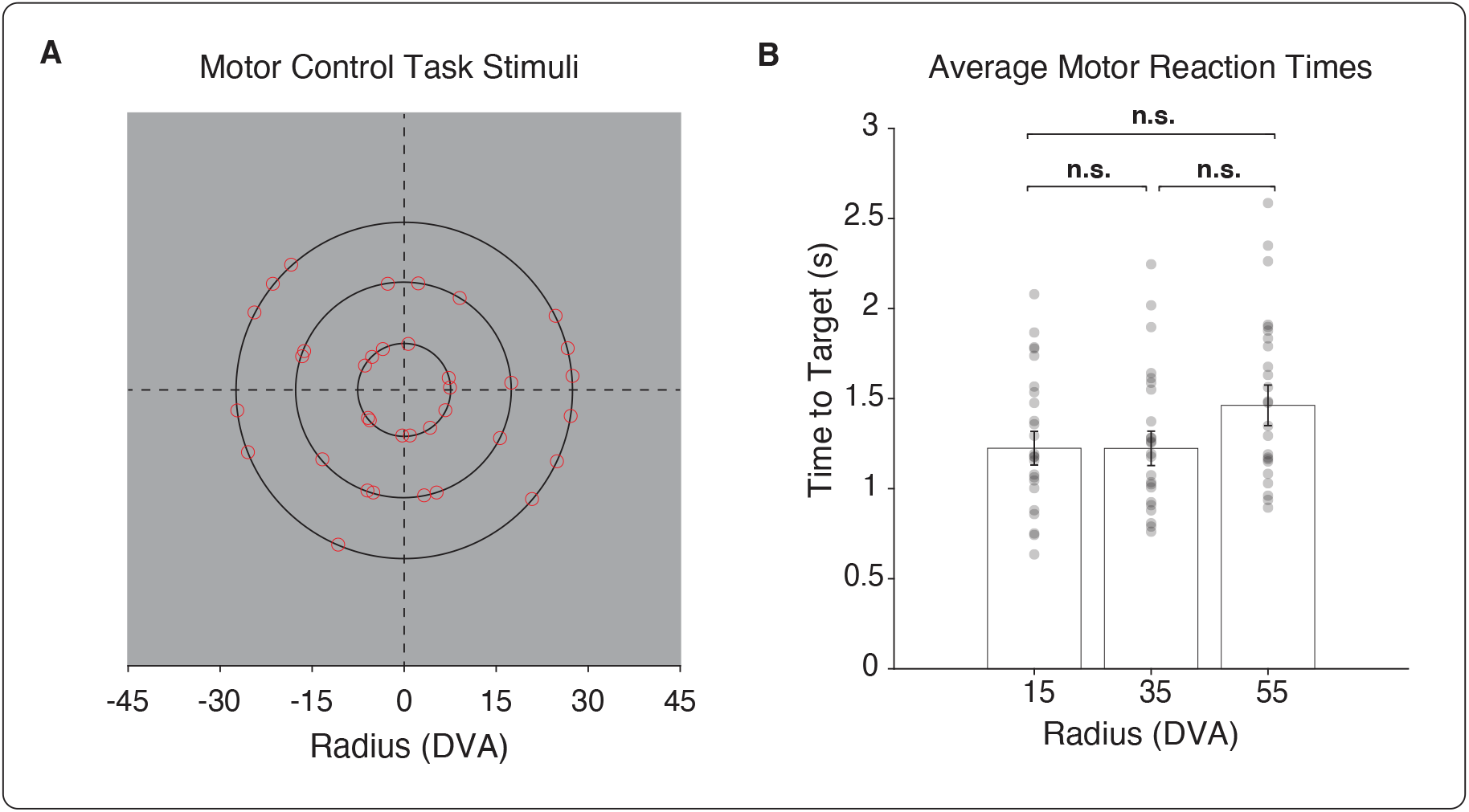
Motor response control trials. **(A)** Participants were asked to orient to a red dot appearing at one of three distances from screen center. **(B)** No differences were found in reaction times binned by distance from screen center (*F*_(2,72)_ = 1.94, *p* = 0.151). Error bars represent 1 SEM. n.s. *p* > 0.05 difference between conditions.

